# Layer-by-Layer Nanoparticle Outer Polyion Impacts Protein Corona Formation

**DOI:** 10.1101/2025.08.13.670086

**Authors:** Tamara G. Dacoba, Simone A. Douglas-Green, Bhuvna Murthy, Alfonso D. Restrepo, Zavian Strom, Margaret Billingsley, Mae Pryor, Paula T. Hammond

## Abstract

Nanoparticles (NPs) can be engineered to achieve targeted delivery with strategies based on surface modifications. These include layer-by-layer (LbL) NPs, modular electrostatically assembled carriers with tunable surface properties altered by changes to the outer polyion layer. Variations in these polymers dictate intracellular trafficking and biodistribution patterns. As NPs are administered, a layer of protein adsorbs to their surfaces, forming a protein corona that affects NP properties, alters biodistribution, and ultimately, impacts therapeutic efficacy. We hypothesized that some differences in LbL NP performance are due, in part, to variations in the resulting protein coronas. To study them, we first optimized an ultrafiltration method to effectively isolate LbL NPs with their protein corona. Following incubation in conditioned media, anionic homopolypeptide outer layers, such as poly-L-aspartic acid (PLD) and poly-L-glutamic acid (PLE), and LbL NPs with the bioinert polymer poly(acrylic acid) (PAA) had the lowest amount of protein associated, lower than conventional PEG liposomes. While mass spectroscopy revealed changes in the protein composition among LbL NPs; albumin, alpha-2-macroglobulin, and apolipoprotein B were most abundant. *In vitro*, pre-formed protein coronas reduced uptake in macrophages but increased uptake in ovarian cancer cells for certain LbL NP outer layers. *In vivo*, LbL NP outer layer influenced both serum half-life and biodistribution. Overall, this work highlights that LbL NPs can be designed to control protein corona formation, and supports that further understanding NP interactions with biological fluids is essential for designing clinically translatable NP platforms.

**Graphical abstract:** 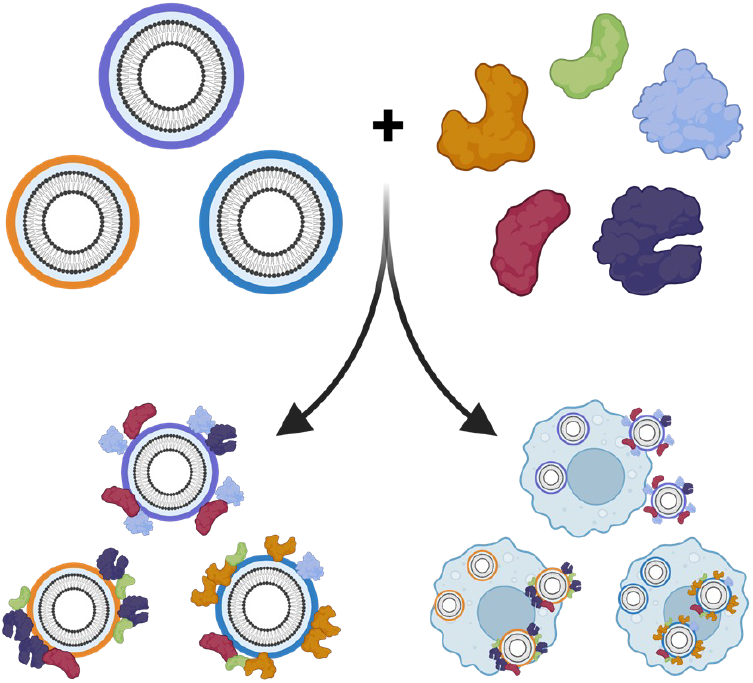

## Introduction

Drug delivery strategies using nanoparticles (NPs) have significantly improved therapies by reducing off-target effects, increasing drug stability, and precisely delivering drug cargo to tissues, organs, or cells of interest [1–3]. These strategies include NPs with varying compositions, structures, and properties, from lipid NPs (LNPs) used for gene therapies to polymeric systems based on poly(lactic-co-glycolic) acid (PLGA) for small molecules to liposomes delivering chemotherapy [1–3]. Across these varied NP platforms, many strategies aim to improve the specificity or potency of NPs by altering their surfaces—including PEGylation to confer stealth or conjugation of targeting moieties to alter distribution. Layer-by-layer (LbL) assembly is an engineering strategy that allows for tuning of the surface properties NPs through the deposition of charged polymers [4–6]. Polyelectrolytes are layered onto an oppositely charged NP core by precise tuning of the ionic strength and pH, and subsequent layers of polymers can be added to form colloidally stable LbL NPs [7]. For these LbL NPs, changes in the final surface polyelectrolyte, also termed the outer layer – which is typically a negatively charged polymer (**Figure 1A**) – can yield different NP biodistribution patterns across healthy and diseased animal models, with some polymers conferring specific receptor affinity. In particular, hyaluronic acid (HA), a well-known CD44 agonist, has been utilized on the NP surface to target tumors overexpressing CD44, such as breast and ovarian cancer [8–10]. Homopolypeptides such as poly-L-aspartic acid (PLD) and poly-L-glutamic acid (PLE) have demonstrated an inherent targeting capacity toward murine and human ovarian cancer cells [11], with recent studies associating this targeting capacity with the presence of amino acid transporters on cancer cells [12]. Conversely, the lack of known receptors for poly(acrylic) acid (PAA) has been leveraged to shield interactions of LbL NPs with circulating granulocytes and monocytes [13]. Similarly, dextran sulfate (DXS) NPs have shown limited targeting of cancer cells, while improving uptake by macrophages and dendritic cells via scavenger receptor [11,14,15]. Beyond known receptor-mediated interactions, the outer polymer layer of LbL NPs has demonstrated impact on biodistribution; for example, HA LbL NPs favor brain accumulation following intravenous administration and PLE-PEG LbL NPs improve the capacity of LbL NPs to diffuse throughout the brain while targeting glioblastoma [16,17]. In all, LbL surface modifications to NP systems have demonstrated broad impact on drug delivery.

**Figure 1.**
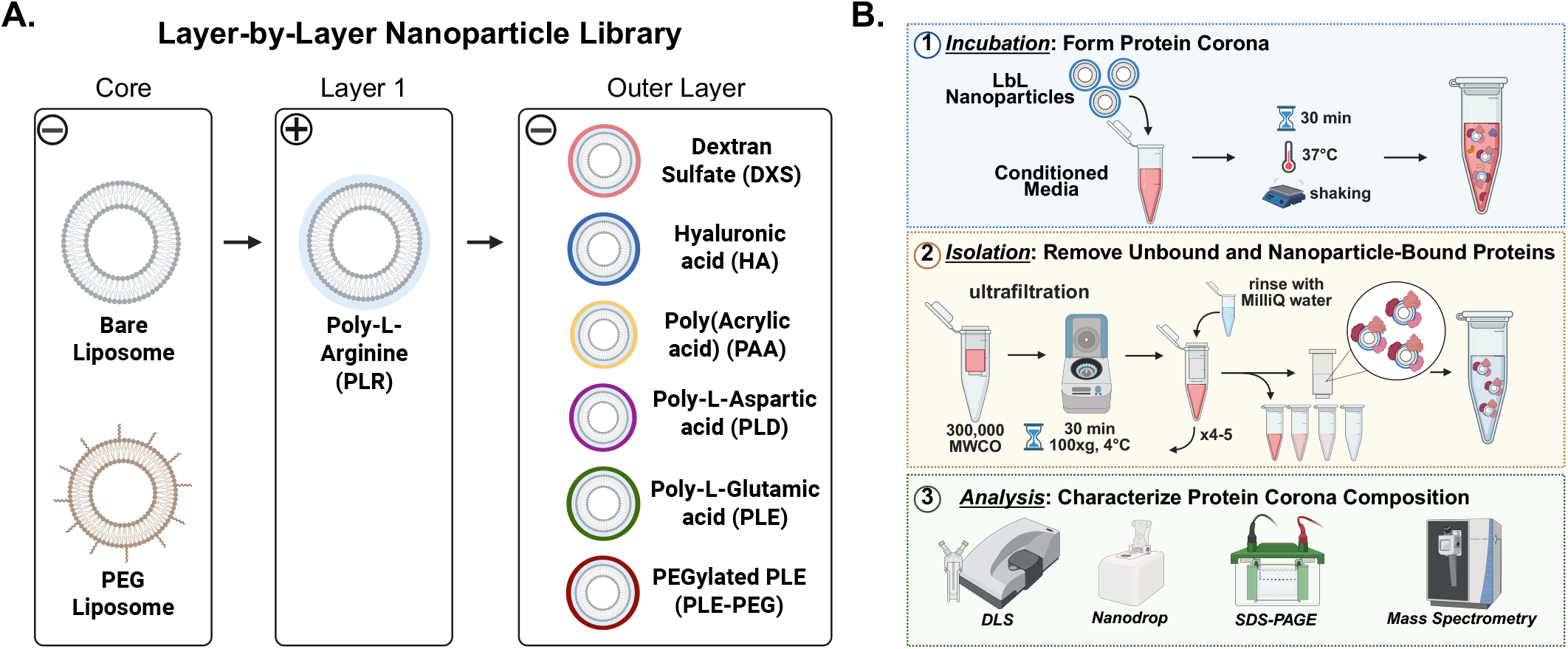
(A) Layer-by-layer nanoparticle (LbL NP) library. Negatively charged bare liposomes were layered with poly-L-arginine, resulting in positively charged PLR NPs. These were further layered with a negatively charged polyelectrolyte (dextran sulfate, hyaluronic acid, poly(acrylic acid), poly-L-aspartic acid, poly-L-glutamic acid, and PEGylated poly-L-glutamic acid) to form the outer layer. PEGylated liposomes were used as a clinically relevant standard. **(B)** Optimized ultrafiltration method to isolate protein-bound LbL NPs from free proteins. (1) LbL NPs were first incubated in conditioned media for 30 min at 37 °C to form a stable protein corona. (2) Then, protein-bound LbL NPs were isolated by ultrafiltration (with a MWCO of 300,000 kDa), for 30 min, at 100 xg, and 4 °C. After several washes, LbL NPs with protein corona were isolated. (3) Last, the isolated protein corona was characterized, including the effects on physicochemical properties of LbL NPs (via DLS), protein concentration (nanodrop), separation of proteins based on molecular weight (SDS-PAGE), and protein composition (with mass spectrometry).

Though surface modification is a promising strategy for targeting NP delivery, the NP surface is immediately altered when it reaches biological fluid. Specifically, proteins and other biomolecules adsorb to the NP surface, forming the so-called protein corona, that directly influences a NP’s *in vivo* behavior. The importance of this alteration is highlighted in 2022 FDA guidelines, reporting that, once administered, nanomaterials can develop different biological properties that affect their safety and efficacy [18,19]. Despite being a critical aspect of NP design and therapeutic effect, the study of the protein corona and its effects on NP performance is still limited [20,21]. Most of the characterization has been carried out for metallic NPs that have more limited therapeutic applications [21]. Given the rise in FDA-approvals for lipid-based NP systems [22], more recent studies have explored the effects of protein corona in lipid nanoparticles (LNPs). For instance, the association of ApoE to LNPs is linked to high liver tropism mediated by hepatocytes [23]. The modification of LNP lipid composition has resulted in different biodistribution patterns [24–26] and proteins adsorbed, with greater amounts of vitronectin associated to lung-tropic LNPs and beta2-glycoprotein for spleen-tropic LNPs [27]. Additionally, albumin, the most abundant protein in blood, has been associated with long circulation times and decreased liver uptake [28].

Based on these correlations between NP composition, protein corona, and biodistribution, we hypothesized that the proteins associated with LbL NPs may influence biodistribution and efficacy *in vivo*. Further, LbL NPs provide an ideal system to observe the impact of NP surface properties on protein corona formation as the modularity of LbL NPs allows for the NP core to be held constant while altering surface polyelectrolyte composition. In this work, we first develop a method to effectively isolate LbL NP-bound protein from free or unbound proteins. Then, using this method, we characterize how protein adsorption changes the physicochemical properties of a library of eight LbL NPs and control PEG liposomes, including changes in NP size and surface charge, as well as the amount of protein associated to the NP surface. The composition of these protein coronas is characterized via mass spectroscopy to determine which proteins or protein families associate with specific LbL NP outer layers. From a biological perspective, we investigate the effect of protein adsorption in the interaction of these LbL NPs with relevant murine cells by pre-forming a protein corona on the NPs before observing NP-cell interaction. Finally, *in vivo* studies show different serum half-life and biodistribution for each of the polyelectrolyte outer layers, which can be partially attributed to the adsorption of different proteins such as albumin, alpha-2-macroglobulin, and apolipoprotein B.

## Materials and Methods

### Materials

Cholesterol, 1,2-distearoyl-sn-glycero-3-phospho-(1’-rac-glycerol) sodium salt (DSPG), 1,2-distearoyl-sn-glycero-3-phosphoethanolamine (DSPE), 1,2-distearoyl-sn-glycero-3-phosphocholine (DSPC), DSPE-N-(Cyanine 5) (DSPE-Cy5), and DSPE-N-[carbonyl-amino(polyethylene glycol)-2000] (DSPE-PEG2000) were purchased from Avanti (Alabama, US). Poly-L-arginine hydrochloride (19 kDa), poly-L-aspartic acid (14 kDa), poly-L-glutamic acid (15 kDa), [methoxy-poly(ethylene glycol)-5000]-block-poly(L-glutamic acid) (20 kDa) were purchased from Alamanda Polymers (Alabama, US). Hyaluronic acid (20 kDa) was purchased from LifeCore Biomedical (Minnesota, US). Polyacrylic acid (15 kDa) and Dextran Sulfate (15 kDa) was purchased from Sigma-Aldrich (St. Louis, US).

The 8-Chamber Lab-Tek II Chambered #1.5 German Coverglass system, wheat germ agglutinin Alexa Fluor™ 555 and 647, and Hoescht 33342 trihydrochloride trihydrate were purchased from Invitrogen (Massachusetts, US). Hank’s balanced salt solution (HBSS) was obtained from Gibco (Montana, US), and 16% methanol-free formaldehyde from Thermo Fisher Scientific (Massachusetts, US).

### Liposome synthesis

Liposomes were prepared as previously described [4]. Briefly, cholesterol and lipid stocks were made in only chloroform, or a chloroform: methanol: water [65:35:8 ratio] mixture, and combined in a flask at a mol ratio of 33 DSPC: 33 cholesterol: 33 DSPG: 0.4 DSPE-Cy5. For PEG liposomes, a mol ratio of 55.7 DSPC: 5.6 DSPE: 33.3 cholesterol: 0.4 DSPE-Cy5: 5 DSPE-PEG2000 was used. The lipid solution was dried into a thin film using a BUCHI RotoVap system until dry. A Branson sonicator bath was heated to 65 °C, at which point the flask with the lipid film was partially submerged in the bath. Ultrapure water was added to re-suspend the lipid film to a 1-2 mg lipid/mL solution, under sonication for 3 min. The solution was then transferred to an Avestin LiposoFast LF-50 liposome extruder, connected to a Cole-Parmer Polystat Heated Recirculator Bath to maintain a temperature of 65 °C. The liposomes were extruded through nucleopore membranes by passing the solution through stacked 400 and 200 nm membranes, 100 nm, and 50 nm membranes.

### Layer-by-Layer assembly

NPs were layered with polyelectrolytes as previously described [4]. Briefly, an equal volume of NP core and polyelectrolyte solution was combined under sonication for a few seconds. Varying weight equivalents of polyelectrolyte to liposome core were used based on the polyelectrolyte layer (**Table 1**). polyelectrolyte solutions were prepared in 50 mM HEPES (pH 7.4) and 40 mM NaCl, except for HA, which was prepared in 5 mM HEPES. Freshly layered particles were allowed to incubate at room temperature for 30 min, then purified using tangential flow filtration (TFF).

**Table 1.**
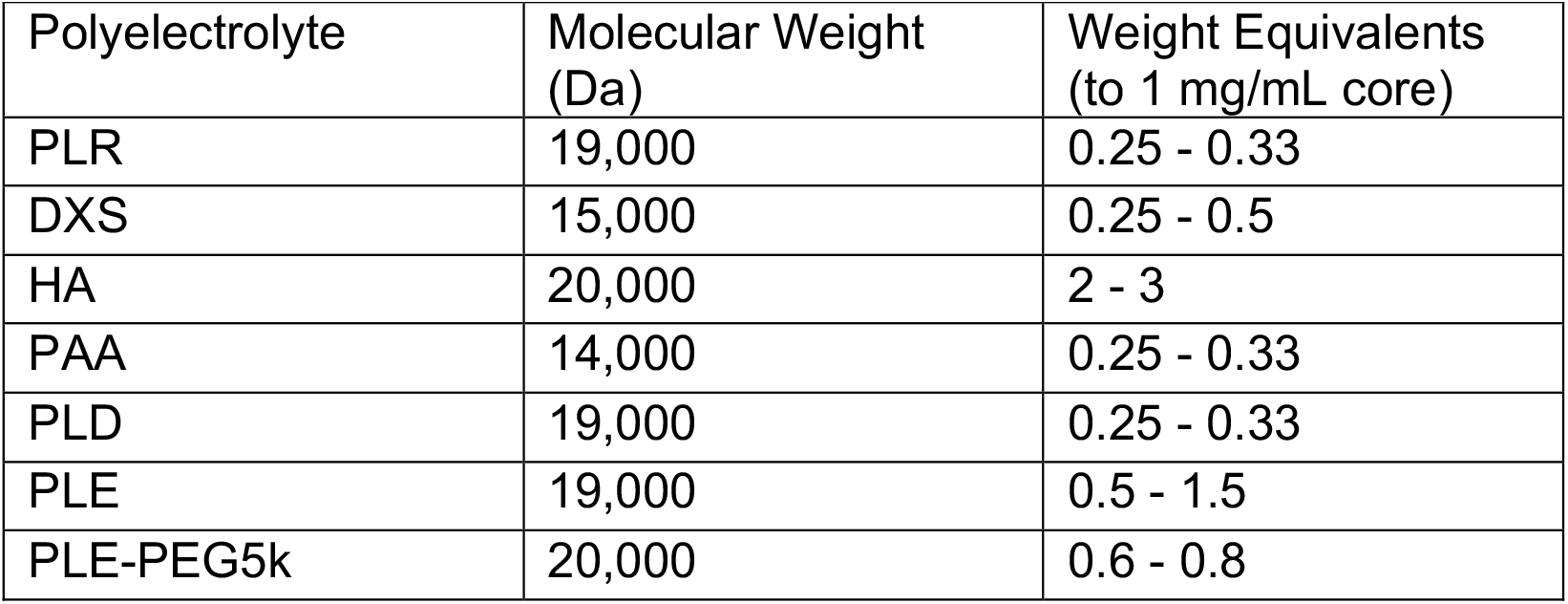
Summary of the polyelectrolytes used for the LbL library with their corresponding molecular weights and the weight equivalents (to a 1 mg/mL lipid core)

### NP Purification by tangential flow filtration

NPs were purified using a Spectrum Labs KrosFlo II filtration system (Repligen, Massachusetts, USA) after each step of the layering process. Hollow fiber filter membranes with 100 or 300 kDa molecular weight cutoffs made of modified polyethylene sulfone were used to remove free lipids or polyelectrolytes from the NP solution. Samples were filtered at 13 mL/min or 80 mL/min (for 13 or 16 tubing, respectively) with an ultrapure water inlet line to replace the volume of waste permeate. NPs were considered purified after collecting at least 5 volume equivalents of waste. Then, the samples were concentrated to the desired volume. After each purification step, NP concentration was tracked based on a calibration curve of the initial liposome solution measuring the Cy5 fluorescent reading (excitation/emission 630/670 nm). For this purpose, NP samples were diluted in a 1:10 ratio in dimethyl sulfoxide and read in a Tecan M1000 microplate reader (Tecan, Männedorf, Switzerland).

### Cell Culture

Murine RAW 264.7 cells were purchased from ATCC and cultured according to manufacturer specifications in Dulbecco’s Modified Eagle’s Medium (DMEM) media supplemented with 10% FBS, 1% penicillin/streptomycin. Murine OV2944-HM-1 (HM-1) cells from Riken BRC were cultured in alpha-MEM media (Gibco) supplemented with 10% FBS, 1% penicillin/streptomycin. RAW 264.7 cells were used for <10 passages from the original stock obtained from ATCC. All cells tested negative for mycoplasma.

### Protein Corona Incubation and Isolation

RAW 264.7 or HM-1 cells were cultured for 2-3 days in their corresponding full media, that was collected and spun down to remove cellular debris. The supernatant, referred to as conditioned media, was aliquoted and kept at –20°C until used in protein corona studies. NPs were incubated in conditioned media for 30 minutes at 37°C while lightly shaking. After incubation, samples were moved to ultracentrifuge filters (Vivaspin 500 Centrifugal Concentrator; 300,000 Da MWCO) and spun at 100 x g for 30 min at 4°C. The permeate, containing unbound proteins, was collected to track protein concentration. The solution remaining in the retentate was washed with cold DI water, which contains nanoparticle-bound proteins, and spun again using the same settings. This process was repeated 4-5 times to ensure removal of free unbound proteins, leaving only the protein corona bound to LbL NPs.

### Nanoparticle characterization

The mean size and polydispersity index (PDI) of the NPs were characterized by dynamic light scattering (DLS), and the zeta potential values by laser doppler anemometry, using a Malvern Zetasizer Pro (Malvern Panalytical, Malvern, UK). The measurements were done in triplicate, at 25 °C, with a red laser (lambda=633nm) and a detection angle of 173°.

For cryo-transmission electron microscopy (Cryo-TEM), 3 uL of the NP sample and buffer containing solution were dropped on a lacey copper grid coated with a continuous carbon film and blotted to remove excess sample without damaging the carbon layer by Gatan Cryo Plunge III. The grid was mounted on a Gatan 626 single tilt cryo-holder equipped in the TEM column. The specimen and holder tip were cooled down by liquid-nitrogen. Imaging on a JEOL 2100 FEG microscope was done using minimum dose method that was essential to avoid sample damage under the electron beam. The microscope was operated at 200 kV and with magnification in the ranges of 10,000–60,000 for assessing particle size and distribution. All images were recorded on a Gatan 2k x 2k UltraScan CCD camera.

### Protein concentration

Protein concentrations were determined based on the protein absorbance at 280 nm for BSA using a NanoDrop 1000 spectrophotometer (Thermo Fisher Scientific, Massachusetts, US). Water was used as blank. For quantification of protein bound to LbL NPs after isolation, the amount of protein retained in the media only control was subtracted from all the NP-containing groups. Permeate of control samples had less than 10% of initial protein.

### Sodium dodecyl sulfate–polyacrylamide gel electrophoresis (SDS-PAGE)

Samples were diluted using 2X or 4X Laemli Buffer (BioRad, California) and normalized by NP concentration. Samples were loaded on a 4–20% Mini-PROTEAN® TGX Stain-Free™ gel (BioRad, California) and run at 120V for 1h in 1X Tris-Glycine-SDS Buffer (BioRad, California). Gels were stained with Coomassie Brilliant blue R-250 (BioRad, California) for 30 min, and destained (10% acetic acid and 10% isopropanol in water) overnight. Gels were then imaged on a ChemiDoc Imaging System (BioRad, California) with the protein gel Coomassie settings to image proteins. Densitometry was done using ImageJ to measure the intensity of the protein bands. Specifically, the 300, 100, 60, and 50 kDa bands were selected; the 300kDa band was normalized to maximum intensity band (corresponding to bare NPs), and the 100, 60, and 50 kDa bands were normalized to the band in the conditioned media stock.

### Mass Spectroscopy

#### Digestion

Digestions were performed with S-trap micro spin columns from Protifi per manufacturer’s protocol, with small variations. Volumes were adjusted for sample volume. First, DTT was added at a 10 mM DTT final concentration, and sample tubes were placed on a heating block for 10 minutes at 95°C. Proteins were then alkylated with iodoacetamide (20 mM final concentration) and incubated for 30 min at room temperature in the dark.

#### LC-MS/MS

The tryptic peptides were separated by reverse phase HPLC (Thermo Ultimate 3000) using a Thermo PepMap RSLC C18 column (2 um tip, 75umx50cm PN# ES903) over a 90-min gradient before nano electrospray using a Orbitrap Exploris 480 mass spectrometer (Thermo). Solvent A was 0.1% formic acid in water and solvent B was 0.1% formic acid in acetonitrile. The gradient conditions were: 1% B (0-10 min at 300 nL/min), 1% B (10-15 min, 300 nL/min to 200 nL/min), 1-7% B (15-20 min, 200 nL/min), 7-25% B (20-54.8 min, 200 nL/min), 25-36 B (54.8-65 min, 200 nL/min), 36-80% B (65-65.5 min, 200 nL/min), 80% B (65.5-70 min, 200 nL/min), 80-1% B (70-70.1 min, 200 nL/min), 1% B (70.1-90 min, 200 nL/min).

The Thermo Orbitrap Exploris 480 mass spectrometer was operated in a data-dependent mode. The parameters for the full scan MS were: resolution of 120,000 across 375-1600 *m/z* and maximum IT 25 ms. The full MS scan was followed by MS/MS for as many precursor ions in a two second cycle with a NCE of 28, dynamic exclusion of 20 s and resolution of 30,000.

#### LC-MS/MS Database Search

Raw mass spectral data files (.raw) were searched using Sequest HT in Proteome Discoverer (Thermo) against a Mouse and Bovine database (Uniprot) and a contaminates database (made in house) with the following search parameters: 10 ppm mass tolerance for precursor ions; 0.02 Da for fragment ion mass tolerance; 2 missed cleavages of trypsin; fixed modification were carbamidomethylation of cysteine, variable modifications were methionine oxidation, methionine loss at the N-terminus of the protein, acetylation of the N-terminus of the protein and also Met-loss plus acetylation of the protein N-terminus. Identified proteins were searched against just Bovine database (Uniprot) and a contaminates database (made in house).

Total spectrum count was reported with accession number and molecular weight. The total percentage of spectrum counts was calculated by dividing the total spectrum count of an individual protein by the sum of spectrum count for each experimental group. UniProt and PantherDB were used to identify the biological functions of identified proteins. ExPASy-Compute pI/Mw tool was used to sort identified proteins by isoelectric point (pI).

### *In vitro* cell uptake

For flow cytometry experiments, 25,000 cells were plated in 100 uL in a 96-well plate and left to adhere overnight. The next day, the media was removed and wells were washed three times with PBS. For the control condition, fresh supplemented media was added, and immediately after, a solution of the LbL NPs in ultrapure water was added as 10% of the well volume. For the pre-formed protein corona condition, NPs were first incubated in conditioned media for 30 min at 37°C, and conditioned media containing NPs (pre-formed protein corona) was then added to the cells. For all these experiments, conditioned media was matched to the cell type evaluated. NPs and cells were incubated for 4 h. After this, cells were transferred to a non-treated 96-well plate, washed with PBS, stained for live/dead with Zombie Violet™ Fixable Viability Kit (BioLegend, California, US). Cells were fixed in 4% formaldehyde and then washed and resuspended in FACS buffer. Cells were read within 4 days of preparation in a flow cytometer BD LSR II or BD Fortessa (BD Biosciences, New Jersey, US). FlowJo™ v10 (BD Biosciences, New Jersey, US) was used to analyze the resulting data. Median fluorescence intensity (MFI) values were normalized to untreated (UTX) cells (MFI_NPs_/MFI_UTX_).

### Confocal Microscopy

Chamber slides were coated with 50 ug/mL of collagen diluted in 0.02N acetic acid, then aspirated, washed with PBS, and let air dry. Chambers were stored at 4 ºC until use. RAW 264.7 or HM-1 cells were plated at a density of 75,000 cells per chamber and cultured overnight. NP dosing was done as described for *in vitro* uptake studies.

After 4h, cells were washed with PBS three times. Cells were then fixed with 4% formaldehyde in PBS for 15 min at room temperature and washed three times with HBSS over 15 min. Samples were stained with 10 µg/mL wheat germ agglutinin labelled with Alexa Fluor™ 555 for 10 min. Then, they were washed with PBS three times over 15 min, fixed with 4% formaldehyde for 2 min, and washed three times with PBS. Cells were stained with 1.5 µg/mL Hoechst 33342 in PBS for 2 min. Finally, samples were washed with PBS and stored at 4 ºC and protected from light until imaging. Imaging was conducted in an Olympus FV1200 Laser Scanning Confocal Microscope with an inverted IX83 microscope equipped with 405, 440, 473, 559, and 635 nm lasers. Images were acquired with 60x objective. Final image analysis was done with FIJI.

### Animal husbandry

All animal experiments were approved by the MIT Committee for Animal Care (protocol 404000660). Healthy male and female C57BL/6J mice (6−8 weeks old) were obtained from The Jackson Laboratory and housed at the Koch Institute animal facility at MIT in groups of up to 5 mice per cage. Animals were allowed to acclimate for at least 72 h before any manipulation. Routine husbandry was provided with the assistance of the MIT Division of Comparative Medicine (DCM) staff. Animals were given ad libitum access to IsoPro RMH 3000 chow (LabDiet) and water and were maintained in 12 h light-dark cycles.

### *In vivo* nanoparticle dosing

Mice were injected retro-orbitally with Cy5-labeled LbL NPs at a lipid concentration of 0.75 mg/mL in 200 uL. All NPs were dosed in sterile-filtered isotonic 5% dextrose. Blood was collected via retro-orbital bleeding in alternating eyes, at 0h, 2h, 4h, 8h, and 24h post administration.

### Blood analysis

Blood was collected in a micro sample tube with clotting activator gel (SARSTEDT Inc., North Carolina, US). Within 1h of the collection, tubes were centrifuged for 5 min at 10,000 x g. Serum was collected, volume raised to 100 uL with PBS, and read in a Tecan M1000 microplate reader (Tecan, Männedorf, Switzerland) to measure the Cy5 fluorescence (excitation/emission 630/670 nm). Values from control mice receiving no NP were subtracted as background signal for all NP groups. Fluorescence values at 0 h were used as the 100% injected dose for each animal group. Half-lives were calculated by fitting the % injected dose to a two-phase decay model, setting the plateau at 0%, and Y0 at 100%.

### Organ harvesting and fluorescence analysis

At 24h after dosing, mice were euthanized and organs were harvested, immersed in RPMI, dried and imaged on a Xenogen IVIS Imaging System (PerkinElmer, Massachusetts). Organ fluorescence was background corrected based on area and values from the same control (untreated) organs. Total recovered signal is calculated from the sum of the total radiant efficiency of all the organs imaged from a mouse. Each organ’s recovered signal is normalized by the total recovered signal per mouse.

### Statistical analysis

Data analysis was performed using GraphPad Prism version 9.1 (GraphPad Inc.). Statistical analysis is described in each figure caption. Data were expressed as the mean ± standard deviation (SD). p values of 0.05 or less were considered statistically significant.

## Results and Discussion

### Ultrafiltration method reliably isolates protein corona on layer-by-layer nanoparticles

To determine the influence of outer layer chemistry on LbL NP protein corona formation, a library of eight LbL NPs with varying outer layers was generated using LbL assembly (**Figure 1A**). PEG liposomes, with 5% PEG, were used as the standard to represent a clinically approved liposomal formulation. All NPs contained a liposomal core fluorescently tagged via a Cy5-phospholipid. The modularity of this platform yielded LbL NPs with very similar physicochemical properties despite the varied polyelectrolyte outer layers, with particle sizes of 120 – 140 nm, polydispersity index (PDI) <0.25, and negative zeta-potential between -40 to -65 mV (except for poly-L-arginine (PLR)– +60 mV) (**Figure S1**). Thus, any differences observed in protein corona formation can be directly associated to the changing nature of the surface properties of the LbL NPs, and not to the variation in other physicochemical properties (i.e., particle size, net surface charge, core composition, mechanical properties) that are known to have an impact in protein adsorption [29–34].

First, to form protein coronas, LbL NPs were incubated at 37 °C, lightly shaking for 30 min in relevant media from macrophages. Previous studies have reported protein corona formation on different types of NPs reaches an equilibrium after 20 min of incubation [35]. As most strategies to isolate NP-bound proteins rely on centrifugation or ultracentrifugation [21,36], we initially attempted to isolate three different LbL NPs (HA, PAA, and PLE NPs) via centrifugation (**Figure S2A**). For this, we incubated the LbL NPs in 10% FBS in water, since this vehicle would enable to more accurately assess the effect of the method and the presence of proteins on LbL NP stability, avoiding confounding influences from the ionic strength of buffers like phosphate buffer saline, which could disrupt the LbL NP assembly. When comparing the differences in size among the LbL NPs before the isolation process to those isolated in water or incubated in 10% FBS, some minimal differences were noted (**Figure S2B**). Nevertheless, the most striking effect was seen on the surface charges. Upon centrifugation of any of the LbL NPs in the water control, a significant loss in the net anionic surface charge could be detected, which was even greater for the LbL NPs incubated in 10% FBS (**Figure S2C**). Given the LbL assembly process is based on electrostatics, the high forces needed to effectively isolate the NPs causes stripping of part of the polyelectrolytes in the outer layer [37]. Therefore, this centrifugation method impacts the surface LbL NPs, making it ineffective for isolating LbL NPs in the study of protein adsorption.

As an alternative, we evaluated an ultrafiltration method with lower centrifugation forces (100xg and 4 °C) using a membrane with a MWCO of 300,000 Da, which would allow for most of the unbound proteins to pass through the membrane, while retaining the LbL NPs and the associated proteins. After several washes, this isolation process for control LbL NPs (in water) did not affect their physicochemical properties, which were comparable to the original NPs (**Figure S3A, B**). However, for the LbL NPs incubated in 10% FBS, a modest decrease in the surface charge absolute value was seen (**Figure S3B**). We also validated that after the five washes, >50% of the LbL NPs were recovered and more than 90% of free protein had been filtered through (**Figure S3C, D**). Therefore, we implemented this method to isolate nanoparticle-bound proteins in the full LbL NP library (**Figure 1B**), with consistent protein filtration and NP recoveries (**Figure S4**).

### Increase in size and variation in surface charge of LbL NPs suggest protein corona formation

Following isolation, we assessed particle size (**Figure 2A**) and surface charge (**Figure 2B**) after incubation in water (control) or in conditioned media from RAW264.7 macrophages (protein corona). Overall, there was little to no change in particle size for PEG liposomes. A modest variation (∼25 nm) could be seen for bare, DXS, PLD, and PLE-PEG NPs. PAA NPs increased their size by ∼80 nm, while PLE, PLR, and HA NPs had an increase in particle size of ≥150 nm. For surface charge, protein adsorption only impacted PLR and HA NPs. PLR NPs charge converted from highly positive to slightly negative, most probably due to the adsorption of negatively charged proteins on the positively charged NPs, which is likely the cause of its size increase as well. In the case of HA NPs, the surface charge of the NPs became more positive – from –40mV in water, to –25mV after incubation with proteins. This modest increase in surface charge is likely affecting the colloidal stability of HA NPs, ultimately causing their aggregation, observed as an increase in particle size. The presence of salts in conditioned media plays a key role in this aggregation, since previously, incubation in 10% FBS in water did not lead to this size increase. Similarly, PLE NPs increased in size, which could be due to the high ionic strength of the media, as previously reported in other biorelevant fluids [16]. Some polyelectrolyte shedding or disassociation from the outermost layer of the LbL NPs could happen upon immersion in the media and contribute to this de-stabilization. Furthermore, cryo-transmission electron microscopy (TEM) images of protein-adsorbed HA NPs or PAA NPs show NP aggregation with proteins bridging several NPs, in comparison to the original LbL NPs (**Figure 2C, D**). Collectively, changes in size and surface charge suggest protein coronas form on LbL NPs with variations between different surface chemistries.

**Figure 2.**
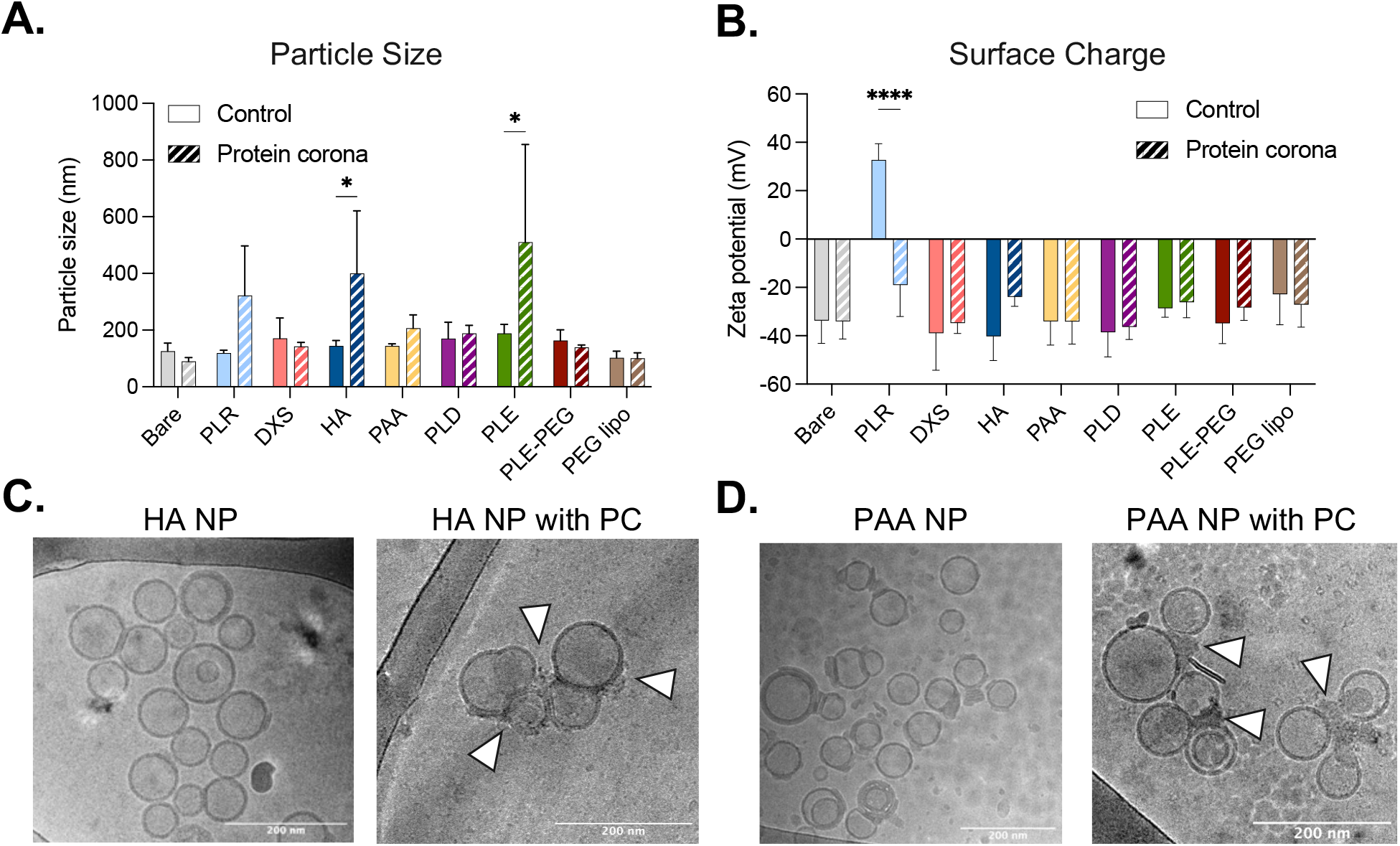
Protein corona formation affects LbL NP size and surface charge. (A) Size (Z-average) and Zeta-potential of LbL NPs isolated in water or after protein corona formation in conditioned media. (B) Cryo-transmission electron microscopy images of (C) HA NPs and (D) PAA NPs in water or after isolation with protein corona (PC). White arrows highlight protein bridging NPs. Scale bars represent 200 nm. Statistical analysis is two-way ANOVA with Šídák’s multiple comparisons test (*p < 0.05, ****p < 0.001).

### Amount of protein corona associated to LbL NPs varies with outer layer chemistry

We further analyzed the amount of proteins associated to LbL NPs, represented as the ratio of protein to NP mass (mg/mg, **Figure 3A**) to account for differences in LbL NP recovery (**Figure S5**). Protein concentration in isolated protein-bound NPs was quantified using a spectrophotometer, while the concentration of LbL NPs was calculated via fluorescent readout (Cy5-phospholipid). HA and PLR NPs presented a much higher association of proteins, with high variability, which aligns with the significant increases in NP size. On the other hand, highly negatively charged homopolypeptides, like PLD and PLE, as well as the bioinert polymer PAA had less associated protein; whereas PEG-containing NPs, either PLE-PEG NPs or PEG liposomes, presented a slightly higher protein association. Similarly, bare and DXS NPs presented comparable protein association values.

**Figure 3.**
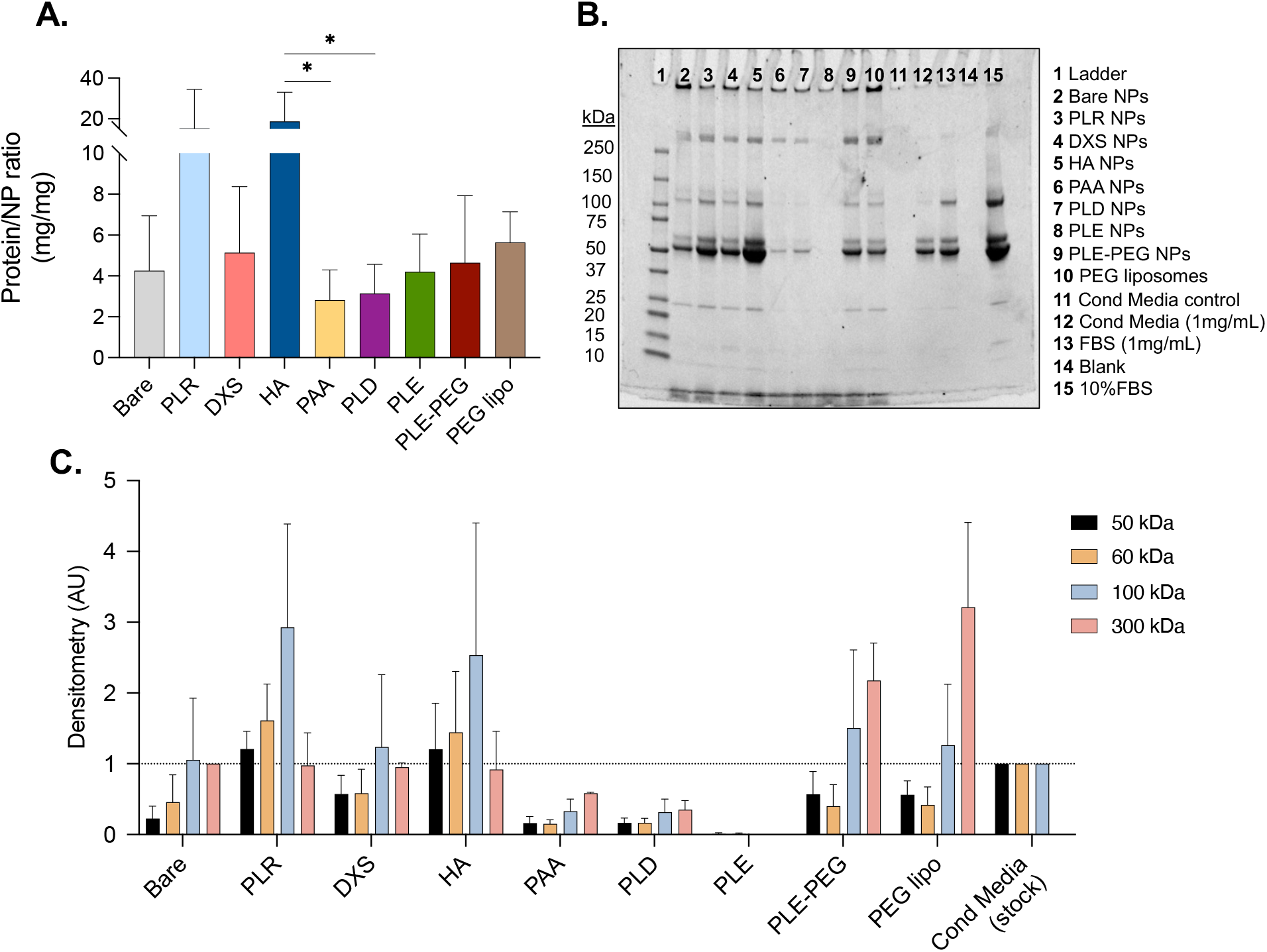
Amount of proteins associated to LbL NPs depends on polyelectrolyte outer layer. (A) Protein to NP ratio (mg of protein per mg of forming lipid) detected on LbL NPs after isolation process. (B) SDS-PAGE of LbL NPs with protein corona. (C) Densitometry analysis of the relative band intensity corresponding to 50, 60, 100, and 300 kDa bands. Statistical analysis is one-way ANOVA with Tukey’s multiple comparisons test (*p < 0.05).

We then ran the samples on SDS-PAGE, normalizing by LbL NP concentration, to identify if any specific banding patterns were detected for the varying outer layer chemistries (**Figure 3B**). We compared the banding patterns of LbL NPs to the ones from conditioned media stock (lane 12) and to the conditioned media control (lane 11)–where the media underwent the same isolation method as all the LbL NPs (Figure 3B). For this, we semi-quantified the bands using densitometry analysis and normalized them to conditioned media stock (for 50, 60, and 100 kDa), or to bare NPs for 300 kDa–due to the absence of a 300 kDa band for the conditioned media control– (**Figure 3C**). In line with the protein to NP ratio, HA and PLR NPs presented more intense bands than any of the other LbL NPs. As expected, the band for albumin (50-70 kDa) was the most abundant one, especially for HA and PLR. Interestingly, the 100 kDa band, corresponding to immunoglobulins, was more evident in the LbL NPs than in the conditioned media alone, suggesting an increased interaction with them. DXS and PEG-containing systems (PLE-PEG NPs and PEG liposomes) presented lower intensity bands for the 50, 60, and 100 kDa bands than the media, but higher than PAA, PLD, and PLE NPs. Notably, PLE NPs showed faint bands across all molecular weights, despite a similar protein-to-NP ratio to PAA and PLD NPs, possibly due to lower staining efficiency of the PLE NP protein composition [38], or an even distribution of proteins near the lower detection limit for SDS-PAGE. Similarly, a stronger intensity band around 300 kDa was seen across LbL NPs, with PLR, PLE-PEG NPs, and PEG liposomes exhibiting the highest values. We cannot fully exclude that this band intensity is a partial artifact of the isolation process, where proteins larger than the MWCO (300,000 Da) might be more retained. Nevertheless, the differences among the different LbL NPs, and the fact that the conditioned media control lane (lane 11) did not show a strong band for these higher molecular weight proteins, suggests this artifact would have a minimal impact. Overall, these results emphasize that outer polyanions such as PAA, PLD, and PLE importantly reduce protein adsorption on LbL NPs, in comparison to traditional PEGylation strategies.

### Varying LbL NP surface chemistry enriches or depletes specific proteins

Differences in the banding patterns and intensities from SDS-PAGE indicate variations in the class of proteins associated with each LbL NP surface chemistry. As such, we sought to identify the types of proteins bound to LbL NPs after isolation using mass spectroscopy. We first looked at the top 12 most abundant proteins found in our conditioned media sample and compared their association with the library of LbL NPs (**Figure 4A**). Overall, changes in the protein corona were influenced by the identity of the outer layer of the NPs. Notably, there is a stark difference between PLE NPs and the other LbL NP outer layers, with PLE NPs having no protein strongly associating with them. Conversely, HA and PAA were the only LbL NPs that demonstrated a strong association with albumin despite it being the most abundant protein in conditioned media. Albumin acts in nutrient and molecule transport and is classified as a dysopsonin, meaning it reduces immune activation and may help improve NP circulation [39]. Further, while all other outer layers associated with apolipoprotein B (ApoB), HA and PLR uniquely presented an increased association with apolipoprotein A-I (ApoA-I). ApoB is known to play a role in lipid transport through VLDL and LDL receptors, including ApoB-100 uptake via LDL receptors in the liver [40,41]. In contrast, ApoA-I has been reported to have a low impact on NP transfection *in vitro* [42], while a recent study leveraged ApoA-I-coated LNPs to favor nucleic acid delivery to hematopoietic progenitor cells [43]. However, there were also notable similarities in the protein coronas across LbL NPs including all outer layers associating with both alpha-2-macroglobulin (A2M) and complement C3. A2M, a protease inhibitor that tags proteases for elimination [44], has been previously linked to uptake via endocytosis by the low-density lipoprotein receptor-related protein [45], while complement C3 is an opsonin with an important role in blood clearance and can induce inflammatory reactions [20]. Interestingly, the already mentioned ApoE that plays a fundamental role in LNP biodistribution toward the liver, was not as highly detected in the protein corona of any LbL NPs. The relative presence of this protein in conditioned media was very low, which could explain these values (**Table S1**).

**Figure 4.**
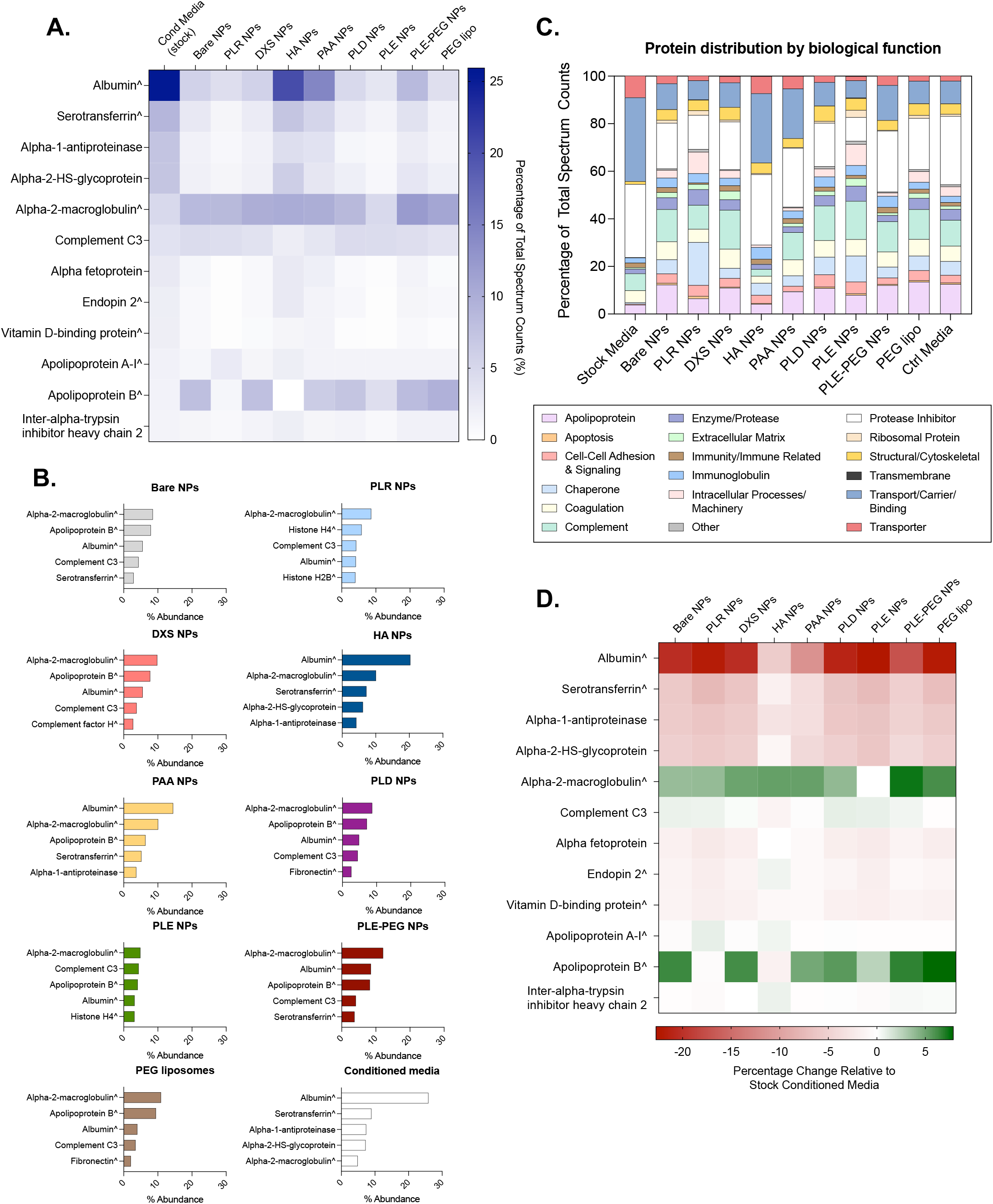
Protein corona composition changes based on outer layer composition of LbL NPs. (A) Percentage of total spectrum counts for the twelve top proteins found in conditioned media from RAW264.7 (stock) and the full LbL NP library. (B) Relative abundance of top hit proteins of each LbL NP. Protein distribution by (C) biological function for the LbL NP library. (D) Enrichment or depletion of top corona proteins relative to stock conditioned media.

Next, for each LbL NP, we identified the five highest associating proteins and analyzed their biological functions, molecular weights, and theoretical isoelectric points (**Figure 4B, Figure S5**). Seven of the LbL NPs had A2M as their top protein, with albumin and ApoB also falling in the top three in most cases (**Figure 4B**). In terms of biological function, most surface chemistries were associated with complement proteins or protease inhibitors, though it should be noted that A2M was classified as a protease inhibitor and may make up a significant portion of that category. The protein corona of HA and PAA NPs contained more transport proteins, likely because albumin was found mostly associated with these outer layers (**Figure 4C**). Across all surface chemistries, more than half of the proteins had a size greater than 70 kDa (**Figure S5A**). Similarly, the isoelectric points of most of the proteins associated with LbL NPs fall within the range of 5 to 6 (**Figure S5B**). With all the LbL NPs being approximately the same size and charge prior to exposure to proteins, these data further corroborate that protein-LbL NP affinities are more likely due to surface chemistries as opposed to other intrinsic NP properties.

We also analyzed how much protein is associated with LbL NP relative to media stock to assess whether proteins were enriched or depleted on the surface of LbL NPs (**Figure 5D)**. While percent abundance allows for a general understanding of which proteins are bound to the NP surface, enrichment and depletion data provide additional insights as to how LbL NPs are interacting with proteins relative to starting abundance in stock, with implications for understanding the protein affinity of various surface chemistries. HA and PAA had the least amount of albumin depleted, further suggesting they have a higher affinity for albumin compared to the other outer layers. We see further evidence of changes in affinities of proteins to various surface chemistries, such as A2M being enriched across all LbL NPs except for PLE NPs, with a similar trend observed for ApoB, that was slightly depleted in HA NPs. All this points to most LbL NPs having a strong affinity toward A2M and ApoB, retaining a high portion of these proteins. Although PEG is typically used to reduce protein adsorption, there is still some enrichment seen for A2M and ApoB in PLE-PEG NPs and PEG liposomes [46,47]. Other relevant studies of NP protein corona formation have identified other proteins such as beta-2-glycoprotein or vitronectin to play important roles in NP biodistribution [27,41]. In our studies, LbL NPs presented less than 0.8% relative amount of these proteins in their protein corona (**Table S1**).

**Figure 5.**
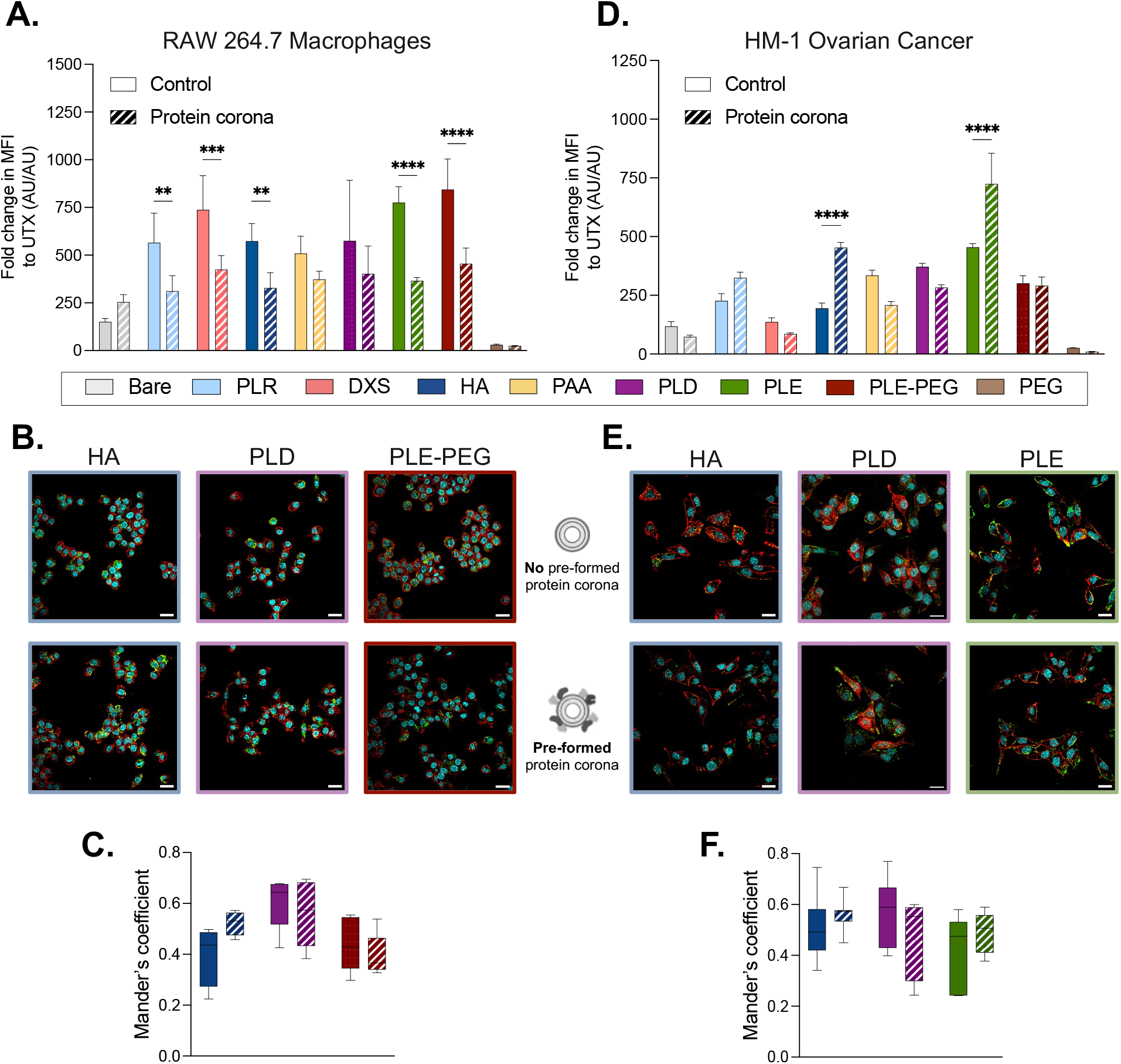
Protein corona formation impacts LbL NP bioactivity *in vitro*. (A) LbL NP association to murine macrophages (RAW 264.7) after 4 h of incubation, with LbL NPs without and with a pre-formed protein corona. (B) Representative confocal images of murine macrophages that were treated with LbL NPs for 4 h, without (top) and with (bottom) a pre-formed protein corona. Images represent the merge of all the color stains. (C) Mander’s coefficient calculation of the NP and membrane co-localization in confocal microscopy. (D) LbL NP association to murine ovarian cancer cells (HM-1) after 4 h of incubation, with LbL NPs without and with a pre-formed protein corona. (E) Representative confocal images of murine macrophages that were treated with LbL NPs for 4 h, without (top) and with (bottom) a pre-formed protein corona. Images represent the merge of all the color stains. (F) Mander’s coefficient calculation of the NP and membrane co-localization in confocal microscopy. Scale bars represent 20 μm. Statistical analysis is two-way ANOVA with Šídák’s multiple comparisons test (*p < 0.05, ***p < 0.005, ****p < 0.001).

### Protein corona affects the uptake of LbL NP in macrophages and ovarian cancer cells

Once we determined that protein adsorption varies significantly based on the surface properties of LbL NPs, we investigated the impact of this protein layer on the interaction with relevant cells. Given the established success of LbL NPs in targeting ovarian cancer cells and immune cells such as macrophages, we chose to assess the impact of protein corona formation on these interactions, testing them in murine RAW 264.7 macrophages and murine HM-1 ovarian cancer cells.

LbL NPs were first incubated for 30 min in conditioned media matched to the cell line to pre-form a protein corona and then incubated with the cells for 4 h. Conditioned media from both cell lines presented a similar protein composition (**Figure S6**. We compared the effect of a pre-formed protein corona with directly adding the LbL NPs to cells in media. Flow cytometry and confocal microscopy were used to evaluate LbL NP association and trafficking, respectively. For macrophages, a generalized decrease in LbL NP association was seen in the case of pre-formed protein coronas, most notably for PLR, DXS, HA, PLE, and PLE-PEG NPs (**Figure 5A**). For control LbL NPs (without a pre-formed protein corona), specific surface polyelectrolytes – such as DXS, PLE-PEG, and PLE –promoted a high NP association with macrophages, aligning with many previous reports showing the increased cell association of LbL NPs. Across groups, the surface deposition of any polyelectrolyte significantly enhanced NP association compared to bare NPs and PEG liposomes. Based on these results, we further investigated whether these variations correlated with changes in trafficking patterns for HA, PLD, and PLE-PEG, aiming to elucidate other changes in LbL NP trafficking due to protein corona formation (**Figure 5B, C**). For HA and PLE-PEG, a pre-formed protein corona slightly altered trafficking *in vitro*, resulting in a subtle higher membrane association. In the case of PLD, the presence or absence of this pre-formed protein corona did not have such a noticeable impact.

In murine ovarian cancer cells (HM-1), pre-formed protein coronas on HA and PLE NPs significantly increased NP association, while it did have a minimal impact for all the other layers (**Figure 5D**). Confocal microscopy on HA, PLD, and PLE NPs confirmed some of these trends (**Figure 5G, H**). For HA, similar trafficking was detected in both conditions, with high association in punctate structures, likely endosomes, via mechanisms previously reported [11]. The pre-formed protein corona on PLD NPs seemed to lead to a lower association with the membrane. The most evident change could be seen for PLE NPs, where protein adsorption decreased PLE NP association to the surface of HM-1 cells, and led to a more even distribution across the cell membrane and cytosol. In this case, protein corona formation of PLE NPs could be affecting their structure and interaction with surface amino acid transporters [12], impacting uptake and trafficking observed.

### In Vivo Studies

Based on the differences in composition of protein corona for each LbL NP, we sought to evaluate if these varied coronas would correlate with *in vivo* circulation times and organ biodistribution. For this, we systemically injected (intravenous, retro-orbital) the seven negatively charged LbL NPs in our library, bare NPs, and PEG NPs, to healthy male and female C57BL/6J mice. We sampled blood during the first 24 h of the study to determine NP serum half-life and evaluated organ biodistribution at 24 h.

We observed that PEG liposomes had the longest fast and slow half-life values (1.91 h and 28.97 h, respectively) among all groups (**Figure 6A, B, Table S2**). Of all the LbL groups, PLD NPs presented the longest circulation half-lives (t_1/2,fast_ 1.21 h and t_1/2,slow_ 11.01 h). Interestingly, the top 5 hit proteins and their relative abundance in PLD NPs protein corona was the most similar to PEG NPs of all LbL NPs, which could explain this trend. However, further analysis must be conducted to thoroughly explore this correlation. HA NPs presented the second longest slow half-life (t_1/2,slow_ 7.10 h), higher than for PAA NPs (t_1/2,slow_ 4.76 h), while their protein corona compositions both included a great relative abundance of albumin (**Figure 4A**). Nevertheless, PAA NPs had much less protein associated with it than HA NPs (**Figure 3A**), which could point to a minimum amount of albumin needed to influence NP half-life. This is further corroborated by previous work in our lab involving HA layering that demonstrated significant increases in NP circulation and retention times in vivo, ranging from 8 to 66 h of slow half-lives [8,48–50]. However, these investigations featured different *in vivo* parameters (whole-blood or whole-body imaging) and/or varied NP cores including quantum dots, polystyrene, or PLGA NPs, which is known to impact cellular interactions [51]. Besides, this past work utilized polyelectrolytes with molecular weights of 40 and 100 kDa, much higher than the 15 to 20 kDa polyelectrolytes used here.

**Figure 6.**
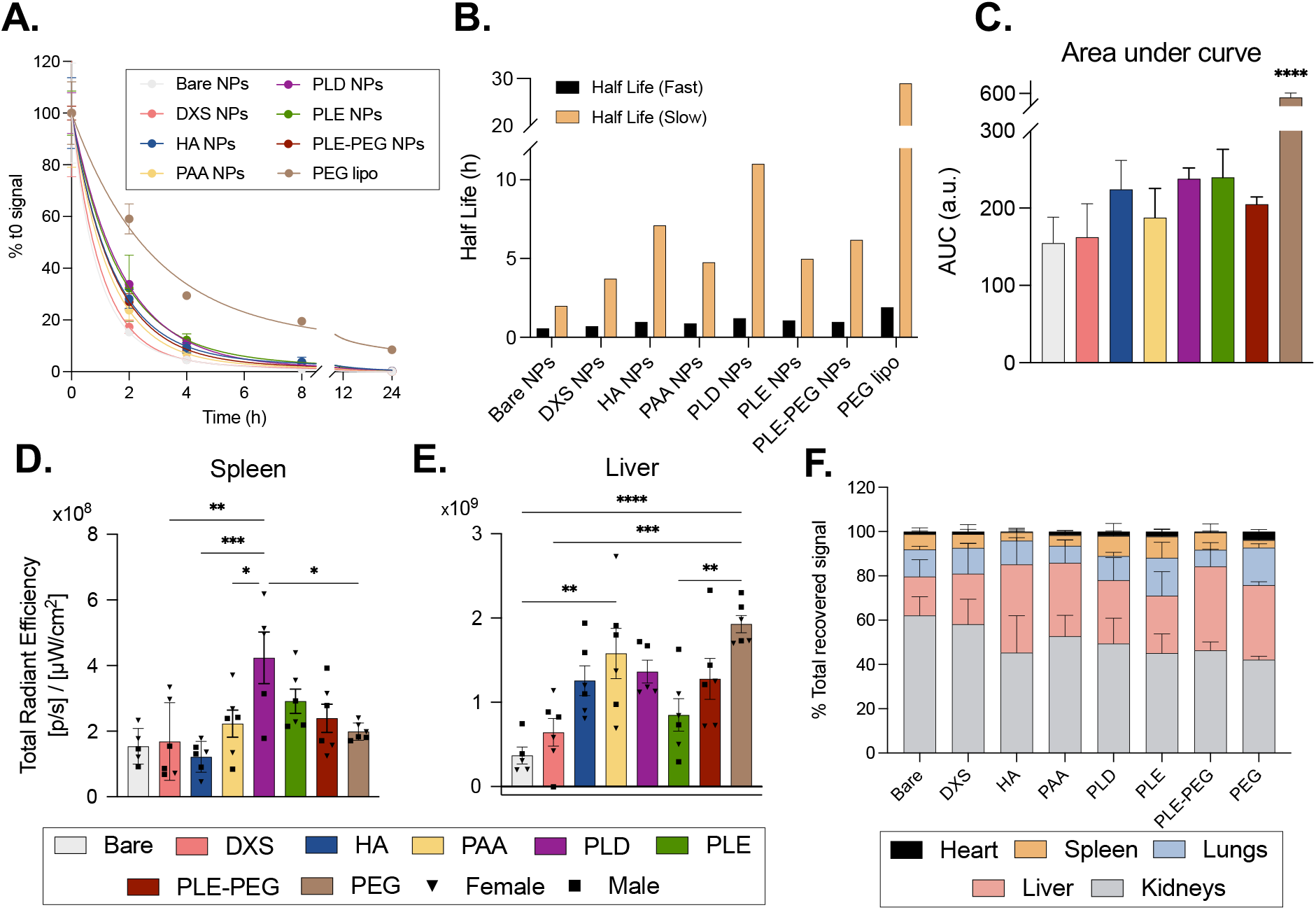
*In vivo* circulation half-life and biodistribution of LbL NPs. (A) Percentage of injected dose of LbL NPs over time and (B) corresponding half-lives fitting a two-phase decay model, after intravenous administration of LbL NPs to C57Bl/6J mice. N=3 male mice per group. (C) Area under de curve for each LbL NP two-phase decay to 24h. (D-F) *Ex vivo* organ biodistribution via IVIS at 24 h following intravenous administration to C57Bl/6J mice, tracking the fluorophore (Cy5)-tagged LbL NPs. Radiant efficiency measurements in (D) liver and, (E) liver. (F) Distribution of LbL NPs in organs based on total recovered signal. Bars represent the mean, and each dot represents one biological replicate: n=3 male (squares) and n=3 female (triangles) mice per group. Statistical analysis is one-way ANOVA with Tukey’s multiple comparisons test (*p < 0.05, **p < 0.01, ***p < 0.005, ****p < 0.001).

The half-lives for both PLE and PLE-PEG NPs differ by ∼1h, suggesting that PEGylation has a modest effect in increasing the half-life of LbL NPs. We did see that PLE-PEG NPs were more stable than the PLE NPs, which might have contributed to this prolonged presence in serum (**Figure 6B**). The overall serum half-life trends across these groups are also evident when comparing the area under the curve (AUC), with PEG NPs having a significantly higher AUC than any of the LbL NPs (**Figure 6C**).

Considering that in these studies we are quantifying the LbL NP presence in serum collected from blood, they omit LbL NPs in circulation that are associating with or internalizing into circulating cells such as monocytes, granulocytes, T or B cells. In fact, a recent study has shown that HA, PLE, and PAA NPs all associate with circulating blood cells to a similar extent but distribute significantly differently across the cell type [13]. Further, as our *in vitro* studies showed that, regardless of outer layer, LbL NPs interact with cells to a greater extent than PEG liposomes (**Figure 5A, D**), we hypothesize that the longer half-life reported for PEG liposomes represents their limited capacity to interact with circulating cells. Thus, the shorter serum circulation time of the LbL NPs may be reflective of cellular interactions in circulation or in organ distribution.

In addition to evaluating the circulation times for these LbL NPs, we compared their biodistribution as protein corona is known to impact organ-level tropism, key to the therapeutic efficacy of many drug delivery vehicles. Thus, we evaluated the organ-level biodistribution of the LbL NPs 24 h after intravenous administration, a timepoint where most of the LbL NPs are outside of blood circulation. First, we looked at the total radiance from each organ. Interestingly, PLD NPs presented a much higher spleen signal than any of the other LbL NPs and PEG liposomes (**Figure 6D**). HA NP signal in this organ was low, in contrast to previous reports in a similar mouse model at shorter time points [13]. This might indicate an early tropism of HA NPs toward the spleen, that decreases over time. Liver accumulation was low for bare, DXS, and PLE NPs (**Figure 6E**). Last, for the heart and the lungs, PEG liposomes presented a significant high signal, likely influenced by the longer circulation times of these NPs (**Figure S7A**,**B**). Similar kidney accumulation was detected across groups (**Figure S7C**). Keeping in mind the important differences in circulation times, that could also bias the total radiance values toward PEG liposomes that had a higher presence in circulation, we further compared the relative organ distribution of each of the LbL NPs (**Figure 6F**). Overall, similar liver and kidney accumulations could be detected across groups which made around 80% of relative organ accumulation for most of the LbL NPs. Bare NPs presented the lowest liver accumulation and the highest kidney accumulation, followed by DXS NPs. Interestingly, HA NPs presented the highest fractional accumulation in the liver, and a low spleen-tropism. In contrast, previous experiments had shown a high spleen signal for HA NPs at shorter time points [13], denoting an early spleen targeting that is lost over time. Both polypeptide outer layers (PLD and PLE NPs), presented a lowered liver accumulation, in favor of accumulation in the heart, spleen, and lungs–in the case of PLE NPs. This higher PLE NP accumulation in the heart and lungs aligns with previous studies at shorter time points [13]. PEG liposomes distributed more toward the heart and lungs. Unexpectedly, PLE-PEG NPs did show one of the lowest heart and lung accumulations, quite opposite to PLE NPs and PEG liposomes, despite similarities in composition. Differences in the stability when comparing PLE and PLE-PEG NPs, the protein corona composition of the three LbL NPs, among other factors, might have led to this difference in relative organ accumulation. Other studies with LNPs had associated the presence of vitronectin with an increased lung accumulation, and of beta-2-glycoprotein for spleen tropism [27]. Nevertheless, these proteins did not stand out as part of the protein corona of these LbL NPs (**Table S1**).

Overall, we have seen that variation in the protein corona across different LbL NPs, together with confounding factors such as the physicochemical properties of the LbL NPs, the NP core, the administration route, and the tissues of interest, have a role in the biological outcomes of LbL NPs. Furthermore, we have focused this study in the role of protein corona formation upon intravenous administration for systemic delivery, while the effect on other administration routes will have to be carefully studied in the corresponding biological fluids [52]. Similarly, it is expected that the composition of the protein corona will change as NPs distribute across other tissues, where new proteins might have a higher affinity [52]. Protein composition in healthy and pathological conditions is also known to change [53], as well as difference based on sex, age and other factors, not taken into consideration in our study [54]. Overall, correlations between surface chemistries and protein coronas, based on protein abundance and amount, warrant further studies in future work, leveraging artificial intelligence and machine learning strategies to discern correlations among these multiple variables [55].

## Conclusion

Overall, understanding how NPs interact with serum proteins is essential for improving targeting and biodistribution, and ultimately, accelerating the clinical translation of nanomedicines. Adsorption of proteins onto NPs once they are administered can affect NP circulation times and delivery properties, including organ-level tropism, cell-level targeting, and intracellular trafficking. Here, LbL NPs enable the isolated characterization of the effect of polyelectrolyte outer layers on protein corona formation. Additionally, the protein corona amount and composition formed around LbL NPs varies depending on outer layer polyelectrolyte, finding albumin, A2M, and ApoB as the most abundant proteins. Through *in vitro* studies, we observed that pre-forming a protein corona modifies LbL NP uptake, in a cell-dependent manner. *In vivo*, we determined differences in slow and fast half-life and biodistribution depending on LbL NP surface chemistry, which may be associated with the presence of proteins such as albumin. Collectively, this work demonstrates the importance of considering the formation of protein coronas on NPs as part of the overall design process, while acknowledging that it is the combination of different inter-related factors in nanoparticle properties that ultimately drives LbL NP efficacy.

## Acknowledgments

This work was supported, in part, by the Bill and Melinda Gates Foundation (grant ID INV-021791 and INV-050202). Research was also funded by the Koch Institute’s Marble Center for Cancer Nanomedicine and NCI Cancer Center Support Grant (P30-CA014051). S.D.G. acknowledges financial support from National Institutes of Health Diversity Supplement (3-R01-EB026344-03S1), Ford Foundation Postdoctoral Fellowship, and Burroughs Wellcome Fund Postdoctoral Enrichment Program (Request ID # 1021694). B.R.M., A.R., and Z.S. were supported by the MIT Undergraduate Research Opportunities Program (UROP). MMB and MP receive support from Break *Through* Cancer. MP receives support from National Science Foundation Graduate Research Fellowship Program. We would like to thank the Koch Institute’s Robert A. Swanson (1969) Biotechnology Center for technical support, specifically the Biopolymers & Proteomics Core, the Flow Cytometry Core, the Microscopy Core, and the Nanotechnology and Nanomaterials core. The schematics of figures (1, 5B, 5E, S2A) were created using https://www.biorender.com. We would additionally like to thank Kimberly Bennett for her help in reviewing this manuscript.

## Competing interests

P.T.H. is a member of the Board of Alector Therapeutics and the Board of Sail Biomedicine, a Flagship company, and a former member of the Scientific Advisory Board of Moderna Therapeutics and the Board of LayerBio. All other authors report no competing interests.

