## Supplemental Information for "Layer-by-Layer Nanoparticle Outer Polyion Impacts Protein Corona Formation"

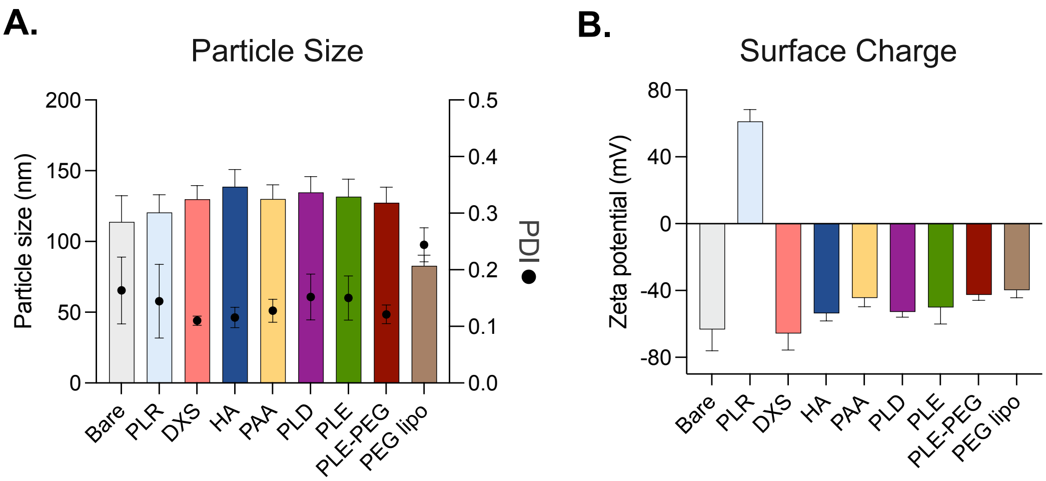


**Figure S1. Physicochemical characterization of the LbL NP library.** (A) Particle size, PDI, and (B) Z-potential of all the LbL NPs and PEG liposomes.

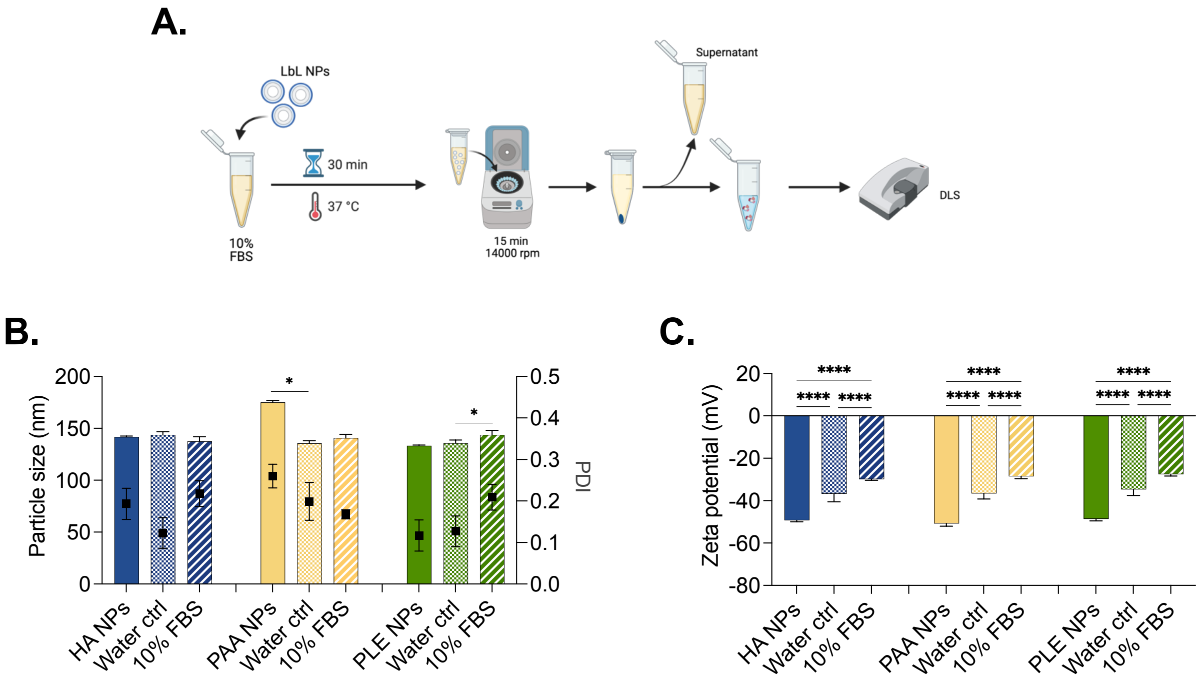


**Figure S2. Validation of centrifugation as an isolation method for protein corona formation on LbL NPs.** (A) LbL NPs (HA, PAA, and PLE NPs) are incubated in 10% FBS in water for 30 min at 37 °C and then centrifuge for 15 min at 14000 rpm. After this time, and pellet os LbL NPs is formed. The supernatant, containing free proteins, is removed. And the LbL NP pellet is resuspended in water for characterization via DLS. Effects of the method on LbL NP (B) particle size and PDI, and (C) zeta-potential comparing LbL NPs before isolation, after isolation of LbL NPs incubated water (water ctrl), or after isolation of LbL NPs incubated with 10% FBS in water.

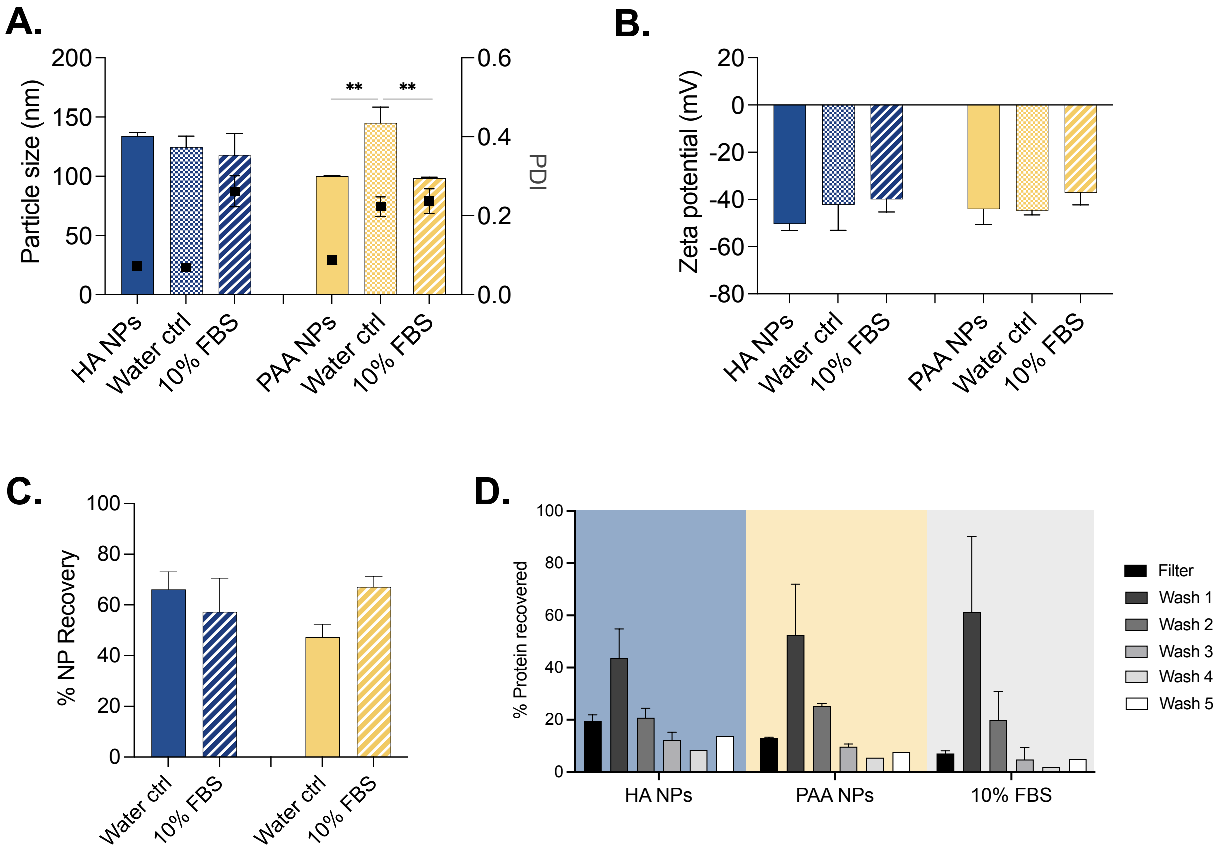


**Figure S3. Initial validation of ultracentrifugation as an isolation method for protein corona formation on LbL NPs.** Effects of the method on LbL NP (A) particle size and PDI, and (B) zeta-potential of the isolation method: comparing NPs before isolation, isolated in water (water ctrl), or after isolation with 10% FBS in water. (C) % of LbL NP recovery after isolation via ultracentrifugation. (D) % of protein recovered in each wash step for LbL NPs and the 10% FBS control.

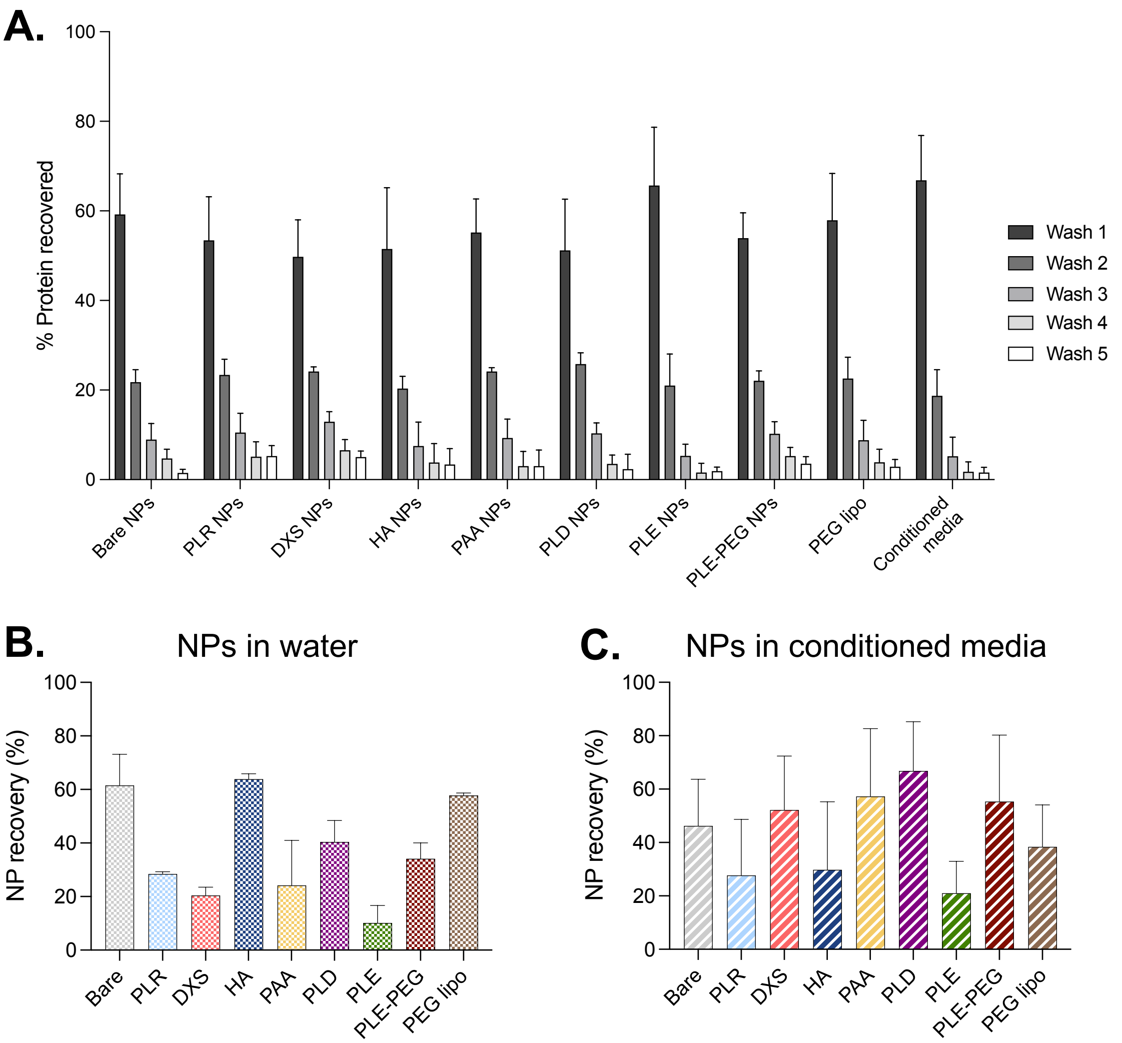


**Figure S4. Protein and NP recovery after ultrafiltration for the full LbL library.** (A) Percentage of protein recovered in each of the washes, and NP recovery (B) in water and (C) in conditioned media.

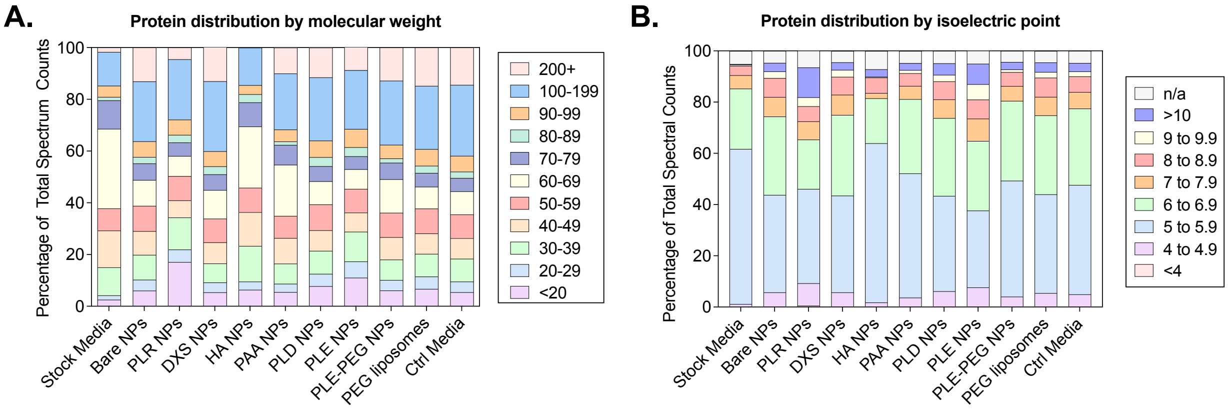


**Figure S5. Protein corona composition changes based on outer layer composition of LbL NPs.** Protein distribution by (A) molecular weight, and (B) isoelectric point for the LbL NP library.

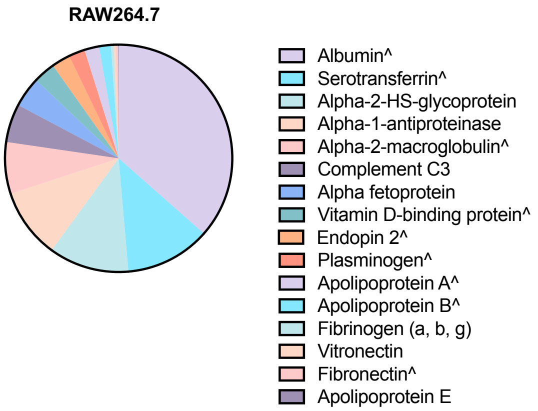


**Figure S6. Relative protein abundance comparing conditioned media from murine macrophages** (RAW264.7, left) **and murine ovarian cancer cels** (HM-1, right).

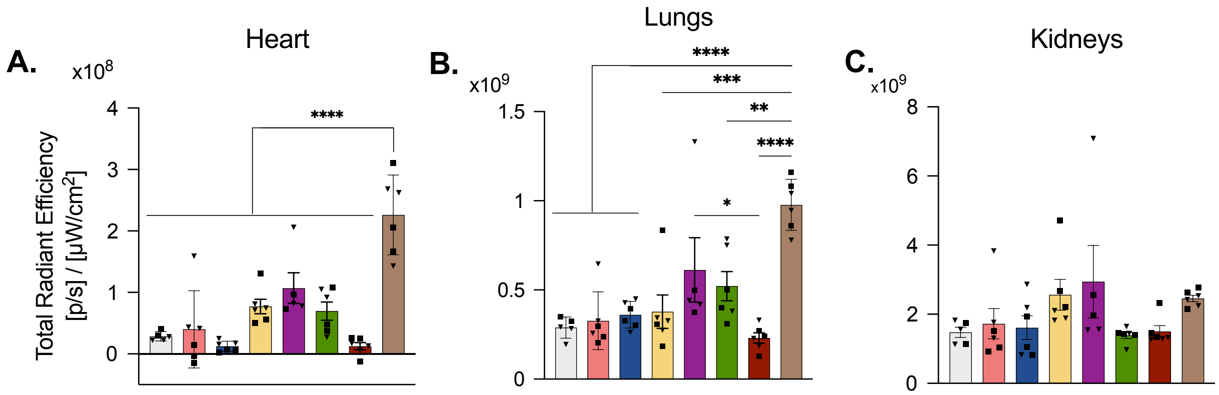


**Figure S7. *Ex vivo* organ biodistribution *via* IVIS at 24 h** following intravenous administration to C57Bl/6J mice, tracking the fluorophore (Cy5)-tagged LbL NPs. Radiant efficiency measurements in (A) heart, (B) lungs, and (C) kidneys. Bars represent the mean, and each dot represents one biological replicate: n=3 male (squares) and n=3 female (triangles) mice per group. Statistical analysis is one-way ANOVA with Tukey's multiple comparisons test (*p < 0.05, **p < 0.01, ***p < 0.005, ****p < 0.001).

**Table S1. Mass spectroscopy data for LbL NPs incubated in conditioned media.** In supplemental spreadsheet.

**Table S2. Fast and slow half-life values of the LbL NP library, after systemic administration to C57BL/6 mice, Values are fit to a 2-decay model, n=3 per layer and time point.**

| | $t_{1/2,fast}$ | $t_{1/2,slow}$ |
|---|---|---|
| **Bare NPs** | 0.57 h | 1.99 h |
| **DXS NPs** | 0.71 h | 3.71 h |
| **HA NPs** | 0.98 h | 7.10 h |
| **PAA NPs** | 0.89 h | 4.76 h |
| **PLD NPs** | 1.21 h | 11.01 h |
| **PLE NPs** | 1.08 h | 4.98 h |
| **PLE-PEG NPs** | 0.98 h | 6.18 h |
| **PEG lipos** | 1.91 h | 28.97 h |
